# Neurodevelopment vs. the immune system: complementary contributions of maternally-inherited gene transcripts and proteins to successful egg development in fish

**DOI:** 10.1101/2021.04.19.440434

**Authors:** Daniel Żarski, Aurelie Le Cam, Thomas Frohlich, Miwako Kösters, Christophe Klopp, Joanna Nynca, Sławomir Ciesielski, Beata Sarosiek, Katarzyna Dryl, Jerome Montfort, Jarosław Król, Pascal Fontaine, Andrzej Ciereszko, Julien Bobe

## Abstract

**Background:** In Metazoans, embryonic development relies on maternally-inherited mRNAs and proteins that are critical for early developmental success and known to play major roles at later stages, beyond zygotic genome activation. However, very poor concordance between transcript and protein levels in oocytes and embryos of vertebrates suggest that maternally-inherited proteins and maternally-inherited mRNAs are playing different roles in unfertilized eggs, not considered to date comprehensively. The aim of this study was to investigate the respective contribution of maternally-inherited mRNAs and maternally-inherited proteins to egg molecular cargo and to its developmental competence using pikeperch, an ecologically and commercially relevant freshwater fish species, as a model.

**Results:** Our data shed new light on the importance of maternally-inherited mRNAs in nervous system development suggesting that neurogenesis is a major mRNA-dependent non-genetic inheritance factor. In contrast, our results highlight a specific role of maternally-inherited proteins in immune response in ovulated eggs suggesting that maternal proteins would rather contribute to developmental success through protection of the embryo against pathogens. Further analysis revealed susceptibility of the transcriptome to modifications during the post-vitellogenic processes (i.e., final oocyte maturation and ovulation), whereas proteomic cargo remains unaffected. This may negatively affect developmental competence of the egg and possibly influence further nervous system development of the embryo.

**Conclusions:** Our study provides novel insights into the understanding of type-specific roles of maternally-inherited molecules in fish. Here we show, for the first time, that transcripts and proteins have distinct, yet complementary, functions in the egg of teleost fish. Maternally-inherited mRNAs would shape embryo neurodevelopment and possibly the future behavior of the fish, while maternally-inherited proteins would rather be responsible for protecting the embryo against pathogens. Additionally, we observed that processes directly preceding ovulation may considerably affect the reproductive success by modifying expression level of genes crucial for proper embryonic development, being novel fish egg quality markers (e.g., *smarca4* or *h3f3a*). These results are of major importance for understanding the influence of external factors on reproductive fitness in both captive and wild-type fish species.

## 1. Introduction

The developmental competence of metazoans ova relies on maternally derived mRNAs (messenger ribonucleic acids) and proteins that drive cellular division until zygotic genome activation (ZGA) [1–3]. Further developmental processes in fish, including cellular differentiation, are followed by reprogramming processes upon ZGA, during which maternally inherited mRNAs undergo degradation and are replaced by zygotic transcripts [4]. In this way, new proteins taking part in embryogenesis are also being produced. However, mRNA degradation upon ZGA is most likely a prolonged process during which selective degradation (targeted to specific mRNAs) occurs [5, 6]. Consequently, it has been found that many mRNAs of maternal origin participate in embryonic development well beyond ZGA, consequently representing a potential nongenetic inheritance pathway [7]; this also probably applies to maternally derived proteins. It is therefore clear that both sets of molecules – transcripts and proteins – of maternal origin are important factors in embryonic development. The lacking or very poor concordance between protein and transcript levels in mammalian oocytes during early development [8] and in X*enopus laevis* eggs and early embryos [9] suggests that maternally derived proteins and mRNAs play different roles during embryogenesis. This hypothesis is additionally supported by a study on zebrafish (*Danio rerio*), in which different functions of transcripts and proteins were suggested in the embryos at 24 h post fertilization [10]. However, this study was performed on embryos at a late stage of embryonic development (during advanced organogenesis, long after ZGA), representing molecular consequences stemming from maternal and paternal factors as well as their interactions. Therefore, there is still a lack of consideration of the specific roles of the two types of molecules of purely maternal origin in unfertilized ova of vertebrates, including fishes. This knowledge is of the highest importance for understanding and identifying the processes and mechanisms responsible for the reproductive capacity of teleost fishes.

In finfishes, oogenesis is the process of oocyte growth, during which proteins and lipids are deposited into the oocyte [11, 12], followed by final oocyte maturation (FOM), during which reorganization of cellular structures occurs [13–15]. This latter process, considered translationally quiescent, is accompanied by transcription within the ovaries, and genomic maturation [12, 16] leads to the deposition of maternally derived molecular cargo in the ovulated, fully mature egg. Upon ovulation, eggs remain transcriptionally and translationally silent or significantly repressed until ZGA [17, 18]. Among the egg constituents, many research efforts have focused on fats and the major fraction of proteins (such as vitellogenins), which serve as nutrients for developing embryos and larvae (as constituents of the yolk sac) [12,19,20]. However, in recent years, increasing attention has been focused on transcripts and the remaining proteins that contribute to embryonic development. There are a number of studies showing that either transcriptomic [1,17,21–24] or proteomic profiles [25, 26] may be used for predicting egg quality and indicating their importance in the developmental competence of eggs. A comparative transcriptomic-proteomic analysis of fully grown ovarian follicles (before ovulation) of zebrafish (*Danio rerio*) identified only a small number (approximately 60) of proteins [27]. Another study characterized the proteome and transcriptome of zebrafish embryos at late developmental stage (i.e., 24 h post fertilization), linking the two types of molecules to crucial developmental processes [10]. However, these studies did not consider the developmental competence of the eggs analyzed and referred to only a single model species. Data on integrated transcriptomic-proteomic analysis from other ecologically and commercially relevant species are still missing. Consequently, a large knowledge gap related to the characterization of the proteome and transcriptome in fully competent ovulated eggs still exists. Additionally, a linkage between transcriptomic-proteomic profiling and the developmental competence of eggs is not available for teleosts. Such information could allow us to identify the processes for which the two types of molecules are responsible and would be indispensable in understanding their role in developing fish embryos. This may further the identification of factors determining egg quality, and more generally factors involved in reproductive biology, in finfishes.

Egg quality in finfishes has serious implications for wild and captive stocks. It has been identified as a crucial factor determining the effectiveness of natural recruitment and commercial production [12,28,29]. Therefore, egg quality – defined as the ability of the oocyte to be fertilized and develop into normal embryos [30] – is attracting much attention from scientists. This allowed us to identify basic extrinsic (such as environmental or nutritional factors) and intrinsic (e.g., *in vivo* aging) determinants of egg quality in finfishes [22,29,31]. However, despite the great progress that has been made, the processes involved in the determination of egg quality are still poorly understood. It should be emphasized that this knowledge is of great importance for understanding the reproductive biology of finfishes, the management of wild stocks and the aquaculture industry [12, 29].

Most of the studies considering egg quality at the molecular level have focused on farmed or model fishes, where a similar approach of categorizing particular samples as either high or low quality has been implemented. This has included the collection of subsamples for molecular analysis immediately after egg collection and monitoring the developmental competence (by determining the embryonic survival rate at various stages of development) of an equivalent subsample during parallel incubation. This approach, although state-of-the-art [29], does not exclude any groups of eggs, a considerable disadvantage. The eggs collected from domesticated fish are of various qualities (including those showing symptoms of overripening and/or intraovarian aging processes), and therefore it is possible that molecular analysis may be performed in cells that are suffering from a lack of basic structural integrity and/or undergoing internal disintegration (also at the molecular level) [24,32–34]. The inclusion of eggs with considerably altered fertilization capacity (which is an important component of egg quality definition [30]), creates the risk of masking many important molecular relationships, limiting the further development of our knowledge. These approaches may also lead to the mixing of unfertilizable eggs with eggs having altered molecular machinery leading to early embryonic lethality – two distinct phenomena. Initial preselection was considered in a study by Żarski et al. [21], where eggs of seabass (*Dicentrarchus labrax*) characterized by apparent abnormalities (as practiced in commercial hatcheries) were discarded from analysis. Additionally, in a recent study on rainbow trout (*Oncorhynchus mykiss*), eggs characterized by overmaturation were excluded from the analysis [17]. These approaches provide new insights into the molecular mechanisms conditioning egg quality, highlighting the good potential of preselecting eggs for molecular analysis. However, in the study of Żarski et al. [21], the strategy undertaken (based on evaluating embryonic survival rate at the 4-cell stage, i.e., 3 h post fertilization) led to the analysis of groups of eggs featuring high variation in quality, which was another limiting factor in such studies. Therefore, a more careful preselection approach based on precisely chosen quality-related indices provides an opportunity to expand our knowledge and would allow us to focus on the molecular mechanisms leading to early lethality (and not related to a fundamental lack of fertilizability), which have been barely considered to date [17].

The aim of this study was to characterize the transcriptomic and proteomic profile of high-quality eggs using pikeperch (*Sander lucioperca*) as a model, the top predator of Holarctic freshwater bodies with high commercial interest [35, 36], and to perform a comparative functional gene set enrichment analysis of the two types of molecules. Next, a comparative analysis of the transcriptomes and proteomes obtained from pikeperch eggs characterized by high and low quality – following a specifically developed preselection procedure – was performed to thoroughly characterize the respective contribution of maternally-inherited mRNAs and maternally-inherited proteins to egg developmental competence.

## 2. Material and methods

### 2.1. Ethics statement

All procedures requiring animal handling (such as reproductive procedures) were performed in compliance with European and national regulations on animal welfare and ethics in experimentation on animals. Whenever appropriate, the procedures were approved by the Lorraine Ethics Committee for Animal Experimentation (CELMEA; APAFIS- 2016022913149909).

### 2.2. General study design

The study comprised of two separate objectives:

1. Objective no. 1: aimed to explore the integrated transcriptomic-proteomic profile of high-quality eggs;
2. Objective no. 2: aimed at comparative transcriptomic-proteomic analysis of eggs characterized by either high or low quality.

Eggs were collected and developmental competence (quality) was evaluated with the same methods in realization of both objectives (described below). Transcriptomic and proteomic analyses were also the same during the realization of the two objectives. Real-time qPCR validation of candidate genes was performed only during realization of objective no. 2. *In silico* and functional gene set enrichment analyses were performed separately for the two scientific objectives undertaken, as specified below.

### 2.3. Broodstock management and reproductive procedures

Pikeperch broodstock (Asialor fish farm, Pierrevillers, France) grown in recirculating aquaculture system (RAS) with fully controlled environmental conditions was used for the study. The RAS was equipped with UV sterilizers, biological filtration and a drum filter to ensure appropriate solids removal. The oxygen levels were kept at >80% saturation, ammonia in the range between 0.2–1.1 mg L^−1^, and nitrite between 0.05–0.70 mg L^-1^. The temperature (within a range of 8–23°C, ±0.5°C) and photoperiod (±1 min) were controlled automatically. The fish had already spawned 3 times before the study was conducted, meaning that they were identified as fully ‘functional’ spawners. To promote the annual gonadal cycle, the fish were exposed to annual fluctuations in the photothermal program described by Fontaine et al. [37] as further modified by Żarski et al. [38] so that the fish were exposed to a wintering period (with temperature below 10°C) for 4 months. Fish were fed throughout the year at a rate of 0.2-1% biomass, depending on the temperature and apparent satiation, with compound feed (50% protein, 11% fat, 10% moisture, 1.55% crude fiber, 1.35% phosphorus, 9.5% ash, and 17.9% nitrogen-free extract; Le Gouessant, France) [38].

Controlled reproduction of the fish was performed according to state-of-the-art protocols. In total, 60 males and 60 females were reproduced following the methods described by Żarski et al. [38]. For induction of ovulation and spermiation, human chorionic gonadotropin (hCG; Chorulon, Intervet, France) was injected into each fish intraperitoneally at doses of 250 and 500 IU kg^-1^ for males and females, respectively. At the time of hormonal treatment, the maturation stage of each fish was scored with the use of catheterization (as described by Żarski et al. [14]). All the fish were identified to be in maturation stage I (representing postvitellogenin oocytes entering the FOM process) out of 6 distinct stages [14]. Next, the fish were periodically monitored for ovulation by gentle pressure of the abdomen. When a particular female was found to have ovulated, the eggs were collected into individual sealable, dry, plastic containers and kept at temperatures between 8-10°C for less than 30 min prior to further procedures. Sperm were collected into dry disposable syringes at between 5 and 7 days following hormonal stimulation (as recommended by Żarski et al. [39]) and were stored at 4°C prior to use (not longer than 15 min), as is practiced in commercial production.

Each collected egg batch was aliquoted into 3 portions. Portion 1 was subportioned into individually labeled cryotubes and immediately snap frozen in liquid nitrogen (later stored at -80°C prior to proteomic and transcriptomic analysis). The second portion was subjected to a preselection procedure (as described further). Portion 3 was subjected to fertilization to assess the egg quality indices.

During the controlled reproduction, the photoperiod was set to 14 h of light per day (provided with neon tube lights), and the temperature was maintained at 12°C from the day of injection until the end of the experiment. Before each manipulation (catheterization, ovulation control, gamete collection, etc.), fish were anesthetized with MS-222 (150 mg L^-1^; Sigma-Aldrich) [40].

### 2.4. Preselection procedure

The preselection procedure is illustrated in Fig. 1. The procedure included placement of the eggs on a plastic Petri dish (to prevent excessive surface adhesiveness of the eggs – a natural feature of pikeperch eggs) and their evaluation under the stereomicroscope for fragmentation of lipid droplets (an indicator of lower quality eggs in percids; [41]) or any other aberrations (symptoms of aging or lack of structural integrity). For the next step, only eggs of normal appearance were used. Next, the eggs were activated with hatchery water, and cortical reaction intensity was evaluated at 3-5 min post water activation, which is a specific quality indicator of pikeperch eggs [42]. The eggs exhibiting at least 90% cortical reaction were further observed for up to 30 min following activation. This was done to determine whether the eggs were able to form blastodiscs at the animal pole (indicating their fertilization capacity – for details, see Żarski et al. [42]), as only the eggs exhibiting blastodisc formation are capable of cellular cleavages following fertilization. The eggs exhibiting a lack of animal poles formation, internal damage or any other irregularities were not considered in further steps of the study (Fig. 1). This preselection procedure allowed us to discard the eggs exhibiting apparent symptoms of lower quality and consider only those eggs with a normal appearance and the potential to be fertilized – an important component of egg quality definition [30]. Consequently, eggs characterized by early lethality were considered to allow us to shed new light on the molecular processes that condition the developmental competence of the eggs.

**Fig. 1.**
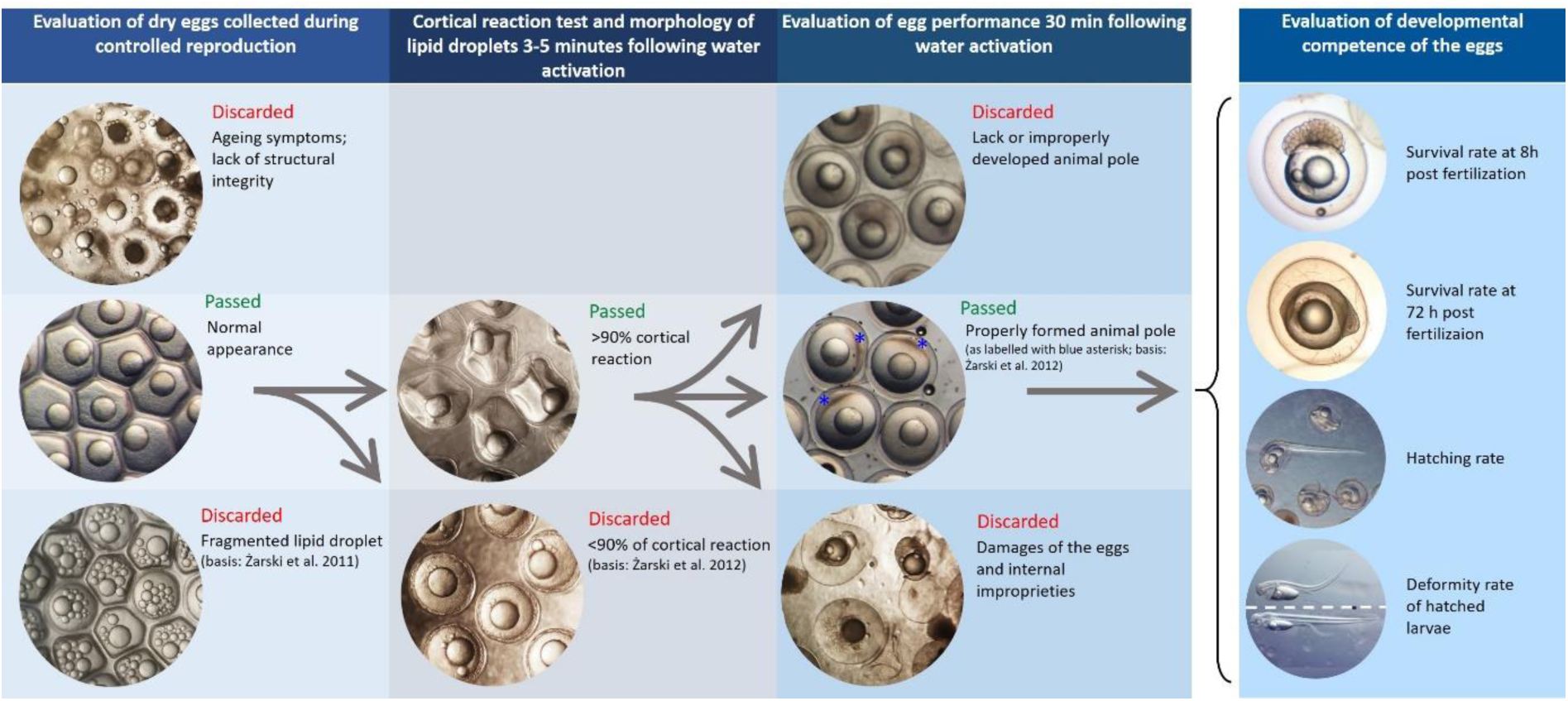
Illustration of the preselection strategy of the pikeperch eggs for the comparative transcriptome and proteome analysis in relation to the developmental competence of the eggs.

### 2.5. Gamete management and fertilization

To assess the egg quality indices, the eggs were subjected to controlled fertilization according to the procedure described by Roche et al. [43]. Briefly, 5 ml of hatchery water was poured over individually labeled glass Petri dishes (30 mm in diameter). Next, a sample of eggs (approximately 100 eggs at one time) was placed onto the Petri dish simultaneously with 50 µl of a sperm sample (pooled sperm obtained from three males each time). Next, the content of the Petri dish was hand-shaken for approximately 10 s to disperse the semen and promote fertilization. Later, the eggs were evenly dispersed across the Petri dish (by gentle hand-mixing) so that they could sediment and adhere to the bottom of the dish (pikeperch eggs exhibit strong adhesiveness to glass surfaces). After 5-10 min, each Petri dish was washed carefully with hatchery water to remove excess sperm and other residues. The eggs were further incubated while adhered to these Petri dishes in an individual 500 ml beaker. The water temperature during incubation was 12°C. Two Petri dishes (duplicates) were established for each egg sample from each female.

For fertilization, freshly collected sperm samples exhibiting at least 80% cell motility (verified under a light microscope at ×400 magnification for each sample as described by Cejko et al. [44]) were always used. The sperm were pooled shortly before use for fertilization, as recommended by Schaefer et al. [45].

#### Evaluation of egg quality indices and designation of samples for molecular analysis

During the incubation period, three indices were recorded: the survival rate (SR) of embryos at 72 h post fertilization (by direct counting of live and dead eggs under a stereomicroscope), hatching rate (HR) of larvae and deformity rate (DR) of hatched larvae. Evaluation of HR was conducted for 5 days (between the 9^th^ and 14^th^ days post fertilization), during which hatched larvae were removed each day from the incubators and counted manually. Next, the larvae were anaesthetized (MS-222, 150 mg L^-1^), and each specimen was individually scored for the occurrence of larval deformity as described previously [46, 47]. DR was calculated as follows: DR = 100 × number of larvae showing deformity/total number of larvae hatched. The analysis was performed with GraphPad Prism 9 software (GraphPad Software, San Diego, CA, USA).

For characterization of the transcriptomic and proteomic profiles, 8 egg batches were characterized by high egg quality (SR > 85%, HR > 60%, and DR < 15%; [47, 48]). For the comparative analysis of high- and low-quality eggs, the five samples characterized by the highest egg quality and the five samples characterized by the lowest egg quality among the preselected samples were used.

### 2.6. RNA extraction

Total RNA was extracted from 50 mg of eggs (approx. 50 eggs) using a Total RNA Mini kit (Cat. No. 031-25; A&A Biotechnology, Gdynia, Poland) following the manufacturer’s protocol. Next, the genomic DNA from the samples was removed using a Clean-Up RNA Concentrator (Cat. No. 039-100C; A&A Biotechnology, Gdynia, Poland) according to the manufacturer’s protocol. The concentration of RNA obtained was measured with a NanoDrop 2000 instrument (Thermo Fisher Scientific, Wilmington, DE, USA). The quality of the RNA was verified with a 2100 Bioanalyzer (Agilent Technologies Inc., Santa Clara, CA, USA). All the samples had RIN (RNA integrity number) values higher than 9.5. The total RNA obtained was aliquoted into two portions and stored at -80°C prior to further analysis. One portion was used for transcriptomic profiling, and the other portion was used for validation of candidate genes with quantitative real-time polymerase chain reaction (qPCR).

### 2.7. Transcriptomic profiling

Transcriptomic profiling of the samples was performed with pikeperch-specific microarrays (8×60k, Agilent), which were designed and successfully validated previously (for details, see Żarski et al. [47]). The microarray design can be accessed via the Gene Expression Omnibus [49] platform under accession number GPL27937. For analysis, the samples were randomly distributed on the microarray slides. Labeling and hybridization of the samples on the microarrays was performed according to the ‘One-Color Microarray-Based Gene Expression Analysis (Low-Input QuickAmp Labeling)’ protocol from the manufacturer (Agilent). Briefly, 150 ng of total RNA from each sample was amplified and labeled using Cy3-CTP. The yield (>1.65 μg cRNA) and specific activity (>9 pmol of Cy3 per μg of cRNA) of the Cy3-cRNA produced were checked with a NanoDrop instrument. From the Cy3-cRNA preparation, 1.65 μg was fragmented and hybridized on a subarray. Hybridization was carried out for 17 h at 65 °C in a rotating hybridization oven. Next, the array was washed and scanned with an Agilent Scanner (Agilent DNA Microarray Scanner, Agilent Technologies, Massy, France) with standard parameters for a 8×60 K gene expression oligoarray (3 μm and 20 bits). The Agilent Feature Extraction software (10.7.1.1) was used to extract the data following the appropriate GE protocol, and the data were further subjected to statistical analysis with GeneSpring GX software (Agilent Technologies, Santa Clara, CA, USA). The gene expression data were scale normalized and log_(2)_ transformed prior to the statistical analysis. Raw data from the comparative analysis of eggs of high and low quality can be accessed via the NCBI Gene Expression Omnibus [49] under the GSE167376 accession number [*the data will be made publicly accessible following publication; reviewers can access the data by using the following token: erkbmaaybjkjpob*].

For the analysis of the transcriptomic profile of n=8 samples of high-quality eggs, only the genes found to be expressed in at least 75% of the samples were used (the gold standard in microarray analysis). Differentially expressed genes (DEGs) between the groups representing high (HQ; n=5) and low (LQ, n=5) egg quality were identified with the use of Gene Spring GX software. The differences between the groups were analyzed by unpaired t-test according to the criteria of a minimum twofold change in expression and a p-value < 0.01. In this analysis, the Benjamini-Hochberg correction was applied. Average linkage clustering analysis (Gene Cluster 3.0) was performed for the differentially abundant genes (unsupervised linkage).

### 2.8. Mass spectrometry-based analysis of egg proteins

Eggs were homogenized for 10 s using a homogenizer (Art-Miccra D-8, Miccra GmbH, Heitersheim, Germany) and by centrifugation at a speed of 23,500 rpm through QIAshredder devices (Qiagen, Hilden, Germany). Samples were then diluted 1:1 with 8 M urea/0.4 M NH_4_HCO_3_, and the protein concentrations were determined with a Bradford assay [50]. Prior to digestion, proteins were reduced for 30 min at 37°C using dithioerythritol at a final concentration of 5 mM and carbamidomethylated with iodoacetamide (final concentration 15 mM) for 30 min at room temperature. Samples of 10 µg total protein were digested for 4 h using 100 ng LysC (FUJIFILM Wako Pure Chemicals, Osaka, Japan). After dilution of the samples with water to 1 M urea, a second overnight digestion step using 200 ng modified porcine trypsin (Promega, Madison, WI, USA) was performed at 37°C. One microgram of peptide was injected into an Ultimate 3000 RSLC chromatography system connected to a Q Exactive HF-X mass spectrometer (Thermo Scientific, Waltham, MA, USA). Samples were transferred to a PepMap 100 C18 trap column (100 µm x 2 cm, 5 µM particles, Thermo Scientific) at a flow rate of 5 µl/min using mobile phase A (0.1% formic acid and 1% acetonitrile in water). For separation, an EASY-Spray column (PepMap RSLC C18, 75 µm x 50 cm, 2 µm particles, Thermo Scientific) at a flow rate of 250 nl/min was used. For the chromatography method, a two-step gradient from 3% mobile phase B (0.1% formic acid in acetonitrile) to 25% B in 160 min and from 25% to 40% B in 10 min was used. Spectra were acquired in the data-dependent mode with a maximum of 15 MS/MS spectra per survey scan.

For data analysis and label-free protein quantification, MaxQuant (v.1.6.1.0) [51] and the *Sander lucioperca* sequences in the NCBI database were used. During the analysis of the proteomic profile of high-quality eggs, only the proteins identified in at least 50% of the samples (i.e., in ≥4 out of 8 samples) were considered abundant and considered in the functional analysis (criteria suggested by Li et al. [52]). Statistical evaluation of the data from comparative analysis of high- and low-quality eggs was performed with Perseus V1.5.3.2. Between-group comparisons were performed with a t-test (followed by permutation-based FDR calculation at a significance level of 5%). The mass spectrometry proteomics data have been deposited in the ProteomeXchange Consortium via the PRIDE [53] partner repository with the dataset identifier PXD023229 [*the data will become freely accessible following publication; reviewers can access the data by logging in to PRIDE* https://www.ebi.ac.uk/pride/ *with the following details: Username; Password: MEXVDYmg*]

### 2.9. In silico, Gene Ontology and functional analysis of transcriptomic and proteomic data obtained during realization of objective no. 1

For both transcriptome and proteome data, protein RefSeq accession numbers for each transcript and protein were obtained. Next, the RefSeq identifiers were indexed against 21,471 human proteins in Swiss-Prot (downloaded from the UniProt database [uniprot.org] on 12 October 2020) with Diamond v0.9.22, followed by alignment of the sequences using BLASTP with an e-value < 10 e^-5^. The resulting file was filtered to keep only the best match for each protein. This enabled us to retrieve gene names and UniProt accession numbers for successfully aligned proteins, which were further used to perform Gene Ontology analysis.

Gene Ontology (GO) and KEGG pathway analyses were performed using the ShinyGO online platform [54] with biological processes considered as targets. Two separate analyses were performed, as presented in Supplementary file 1.

The first GO analysis (Supplementary file 1a) aimed at comparative analysis of the full transcriptome and proteome. To this end, each data set (transcriptome and proteome) was evaluated following the same analytical approach. First, the higher level GO categories were obtained (100 categories for each data set) and compared to each other for the occurrence of common and distinct GO categories. Next, enrichment analysis was performed to obtain 500 significantly enriched (FDR < 0.05) terms for each of the data sets. The obtained terms were further subjected to network analysis based on hierarchical clustering (related GO terms were linked together only when they shared overlap of at least 50% of genes). This allowed us to identify clusters (at least four different ontological terms joined together) characterized further with additional enrichment analysis of the gene list for each cluster, which enabled us to identify biological processes characteristic of each cluster.

The second GO analysis (Supplementary file 1b) was followed by the identification of transcriptome-specific, proteome-specific and conspecific subsets of genes, as revealed by a Venn diagram [55]. Each of the subsets was later analyzed with the ShinyGO platform in three steps:

- Step 1: Enrichment analysis (identification of the 30 most enriched processes with FDR < 0.05) was performed;
- Step 2: Hierarchical clustering was performed to identify separate clusters (sharing at least 30% of genes), with the condition that each cluster include at least three ontological terms;
- Step 3: Additional enrichment analysis of the genes constituting separate clusters was performed to find the 3 most enriched terms representative of each cluster.

Additionally, for each subset of genes, the 5 most enriched KEGG (https://www.kegg.jp/) pathways were identified.

### 2.10. In silico, Gene Ontology and functional analysis of DEGs obtained from realization of objective no. 2

For each DEG, a human homolog was obtained (as described in the previous section). Next, hierarchical clustering-based networking of the 30 most enriched (FDR < 0.05) biological processes was performed with the ShinyGO platform. GO terms were connected to each other only when they shared at least 50% of their genes. In this way, four different clusters connecting at least 3 different GO terms were identified. Next, the genes forming each cluster were once again subjected to GO enrichment analysis, enabling the identification of the 3 most enriched (FDR < 0.05) GO terms characterizing the analyzed DEGs. Additionally, all the DEGs were mapped to clusters of biological processes that were significantly enriched in the transcriptomic and proteomic profiles of high-quality eggs to determine their relevance in whole processes specific to either the transcriptome or proteome.

### 2.11. Designation of candidate gene markers of egg quality in pikeperch during the realization of objective no. 2

To identify the most suitable candidate gene markers to enable the prediction of egg quality in pikeperch, zebrafish gene identifiers for DEGs (identified during comparative analysis of high and low egg quality) were retrieved using ShinyGO. Next, the expression pattern (provided as the number of transcripts recorded per million reads; TPM) during zebrafish embryonic development for each gene was obtained from the Expression Atlas (https://www.ebi.ac.uk/gxa/home) based on the data published by White et al. [56]. Next, the data were grouped into 5 developmental periods (pre-ZGA, blastula/gastrula, somitogenesis, prim stage and 2-5 days post fertilization) identified by White et al. [56] as representative crucial developmental phases. For these periods, the mean expression value was calculated. Furthermore, the data were log_(2)_ transformed and arranged in ascending order based on the pre-ZGA phase as the phase during which the maternal transcriptome plays a major role. The data were visualized as a heatmap with a color scale representing the normalized expression level using GraphPad Prism 9. As the most suitable candidate gene markers, 20 genes characterized with the highest expression during the pre-ZGA phase in zebrafish were considered and taken for qPCR validation.

### 2.12. qPCR validation of candidate gene markers of egg quality in pikeperch during the realization of objective no. 2

Total RNA obtained from eggs characterized by either high or low egg quality was subjected to reverse transcription (1.7 ng of total RNA) using a RevertAid First Strand cDNA Synthesis Kit (Cat. No. K1622, Thermo Fisher Scientific) according to the manufacturer’s protocol. The protocol involved oligo(dT)_18_ primers as well as an optional incubation step (5 min at 65°C) aiming at the removal of secondary structures.

Real-time quantitative polymerase chain reaction (qPCR) was performed for each of the selected candidate genes using a Viia7 (Applied Biosystems) thermocycler. For each qPCR (reaction volume 20 µL), 10 ng cDNA template was used along with DyNAmo HS SYBR Green qPCR Master Mix (Cat. No. F410XL, Thermo Fisher Scientific) and 0.5 µM forward and reverse primers (sequences of all the primers are provided in Supplementary file 2), which were designed with the Primer3Plus online platform [57]. The following cycling conditions were applied: enzyme activation for 10 min at 95°C followed by 40 cycles of denaturation at 95°C for 15 s and annealing and elongation at 60°C for 1 min. After each amplification, melting curve analysis was performed to verify the amplification specificity and compare it with the predicted melting curve verified with uMELT [58]. In the analysis for each gene, a standard curve was calculated using a series of 6 twofold dilutions to determine reaction efficiency (reaction efficiencies between 85 and 110% were considered acceptable). Relative expression for each gene was normalized as the geometric mean of expression values recorded for 4 reference genes, which were chosen on the basis of their stable expression levels and close-to-mean expression values in the microarray analysis [21]. Each reaction for qPCR validation was performed in duplicate. The data were compared between the groups (low and high egg quality) using a t-test (GraphPad Prism 9). Differences between groups were considered significant at p < 0.05.

## 3. Results

### 3.1. Scientific objective no. 1: Transcriptomic-proteomic characterization of eggs

#### 3.1.1. Transcriptomic profile of high-quality eggs

Transcriptomic analysis revealed 10,238 expressed genes encoding unique proteins, including 9,465 proteins for which human homologs could be identified, encoded by 9,233 genes, which were considered in further analysis. The 5 most highly expressed genes, based on the expression level in the microarray, were *hdgfl2*, *cldn4*, galactose-specific lectin nattectin-like (for which no human homolog was found) and *mt-nd5*. All the identified genes are presented in Supplementary file 3.

#### 3.1.2. Proteomic profile of high-quality eggs

The proteomic analysis allowed us to identify 1450 proteins (Supplementary file 4), of which 806 could be detected in at least 50% of the samples and were considered abundant. The most abundant proteins were fish-specific vitellogenins and galactose-specific lectin nattectin-like proteins. For 790 of these 806 abundant proteins, human homologs could be found, which corresponded to 699 unique proteins taken for further Gene Ontology analysis. The most abundant proteins in the eggs were found to be fish-specific proteins, such as various forms of vitellogenins, galactose-specific lectin nattectin-like proteins and ladderlectin-like proteins. The proteins for which human homologs were found to be the most abundant were zona pellucida sperm-binding proteins 3 and 4 (encoded by *zp3* and *zp4*, respectively). All the proteins considered abundant are presented in Supplementary file 5.

#### 3.1.3. Functional analysis of transcriptome and proteome datasets

The transcriptome and proteome datasets were each grouped into 100 higher level Gene Ontology biological process categories (Supplementary file 6). Among 94 conspecific categories, 10 were found to cover various processes linked to reproduction, 6 were related to development, and 12 were related to the immune system (Fig. 2). The remaining categories covered many important but very general biological processes related to, among others, methylation, cell function and maintenance, regulatory processes, protein processing and cell responses to various factors. Among the 6 proteome-specific terms, the acrosome reaction process included only two proteins: zona pellucida 3 and 4, which were among the most abundant proteins. Of the 6 transcriptome-specific higher level Gene Ontology terms identified, three were found to comprise genes involved in various processes affecting behavioral features (i.e., multiorganism behavior, regulation of behavior and intraspecies interaction between organisms) (Supplementary file 6).

**Fig. 2.**
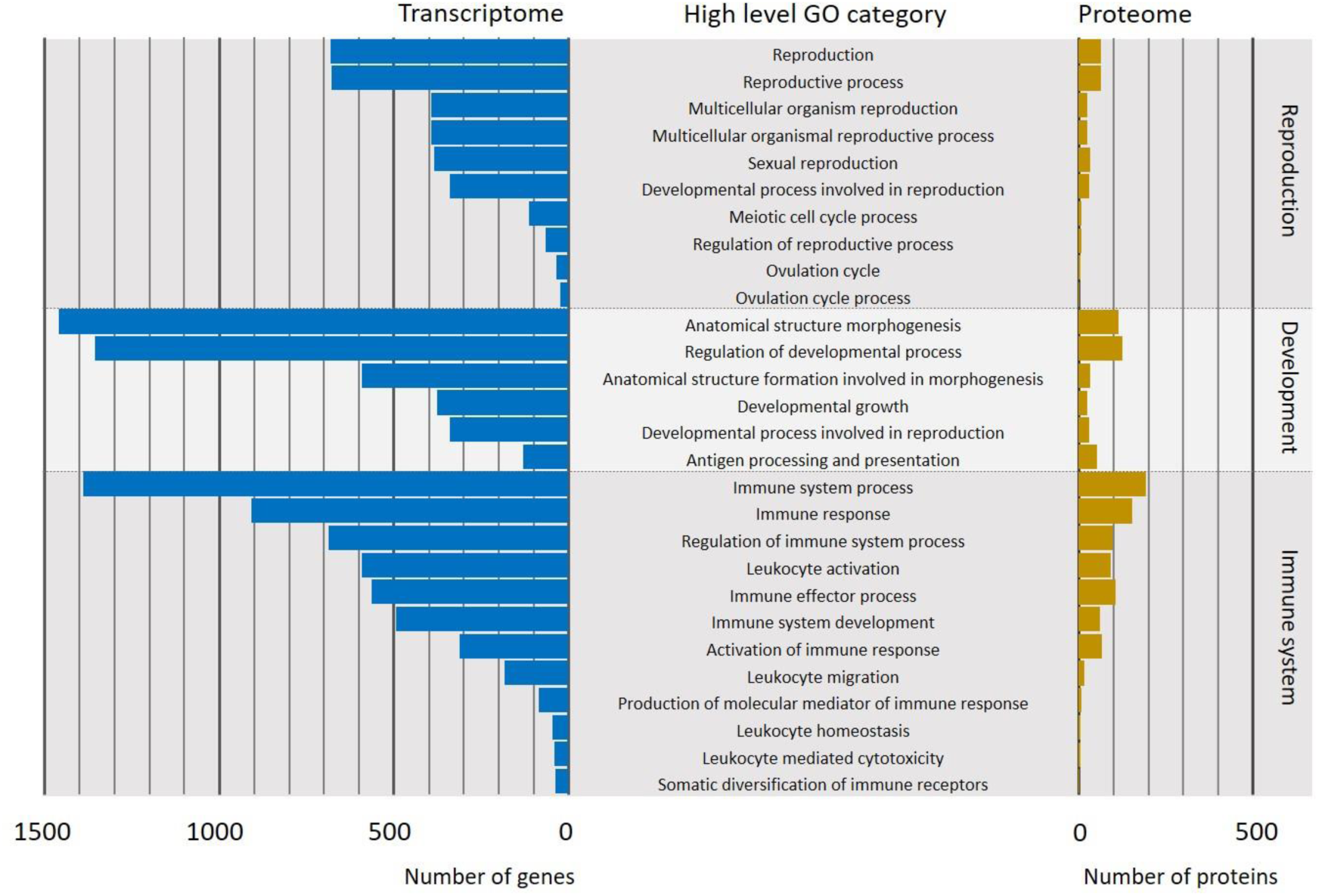
Number of genes and proteins associated with higher level Gene Ontology (GO) categories related to reproduction, development or immune system identified in the transcriptome and proteome of high-quality pikeperch eggs (n=8). For further details, see also Supplementary file 6.

Enrichment analysis yielded the 500 most enriched GO terms for each dataset (i.e., transcriptome and proteome) (Supplementary file 7). Among the transcriptomes, the most enriched (with FDR<3.08e^-09^) processes were clustered into several groups taking part in, among other functions, neurogenesis, mitosis, biosynthesis of molecules (including gene expression), protein modification, intracellular signaling, cell metabolism, transport of molecules and assembly of cellular components (Fig. 3).

**Fig. 3.**
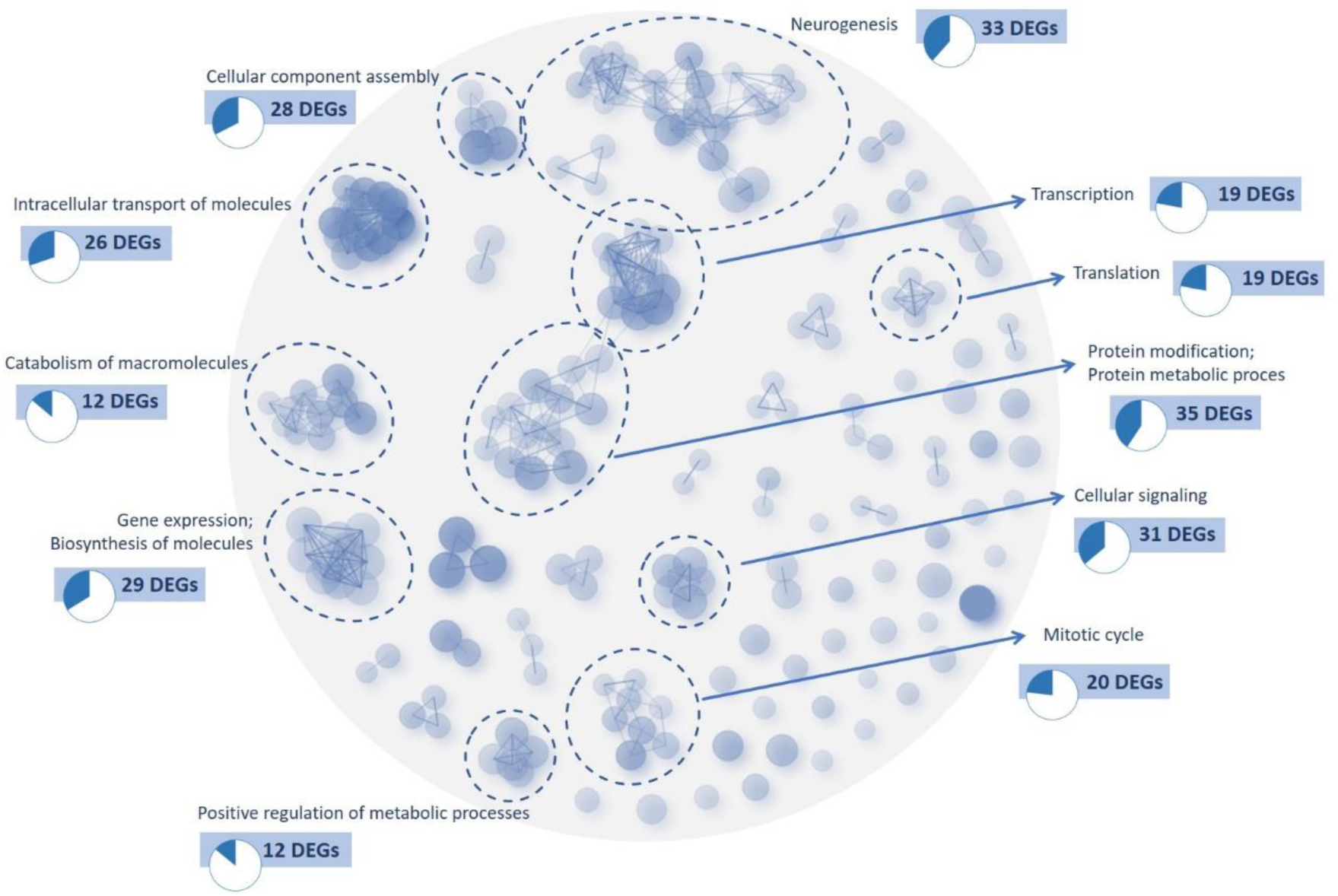
Clustering of the 500 most enriched biological processes (BPs) obtained during the functional enrichment analysis (FDR<0.05) of all the genes identified in the transcriptome obtained from high-quality pikeperch eggs (n=8). For further details, see also Supplementary file 7. Circled clusters are those comprising at least four different BPs, for which additional functional analysis was performed to identify the main BP. Additionally, the number of differentially expressed genes (DEGs) identified between high- and low-quality eggs (for details see also Supplementary file 9) mapped to each of the main BPs were visualized.

The most enriched processes (FDR<2.12e^-05^) identified on the basis of the proteome were clustered into six distinguishable functional categories. These terms included translation, catabolism of proteins and RNA, regulation of gene expression, immune response and biogenesis of cellular components, and cellular metabolic process, with terms related to either metabolism or biosynthesis of purines being highly represented (14 terms) (Fig. 4).

**Fig. 4.**
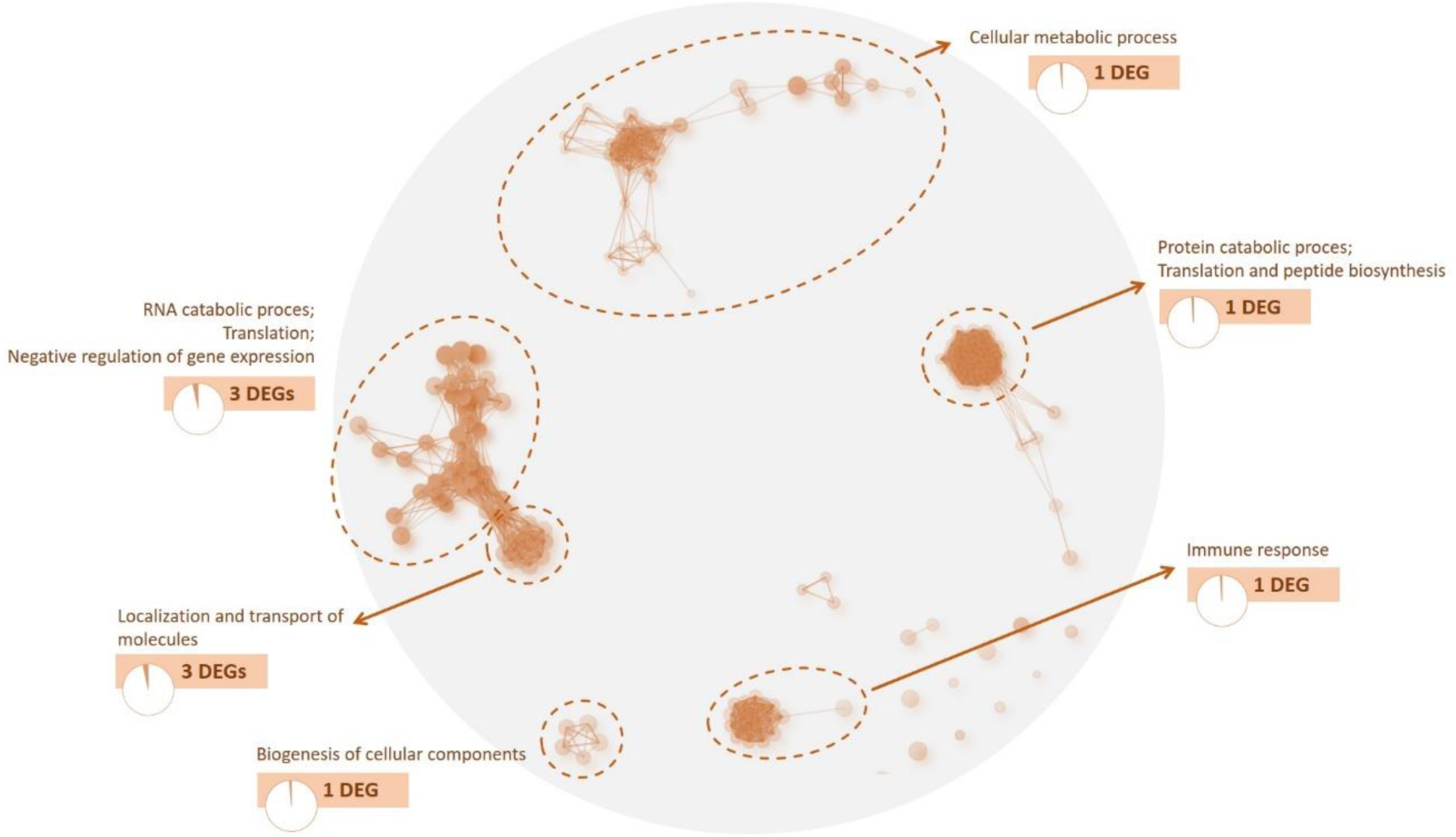
Clustering of the 500 most enriched biological processes (BPs) obtained during the functional enrichment analysis (FDR<0.05) of all the proteins identified in the proteome obtained from high-quality pikeperch eggs (n=8). For further details, see also Supplementary file 7. Circled clusters are those comprising at least four different BPs, for which additional functional analysis was performed to identify the main BP. Additionally, the number of differentially expressed genes (DEGs) identified between high- and low-quality eggs (for further details, see also Supplementary file 9) mapped to each of the main BPs were visualized.

#### 3.1.4. Functional analysis of genes specific to or shared between the transcriptome and proteome

Among all the genes and proteins identified to be present in the high-quality eggs of pikeperch, 509 were found to be conspecific to both data sets. Among all the transcripts identified, 8,724 did not correspond to proteins in the proteome, and among all the proteins identified, 190 were not encoded by genes identified in the transcriptome (Fig. 5).

**Fig. 5.**
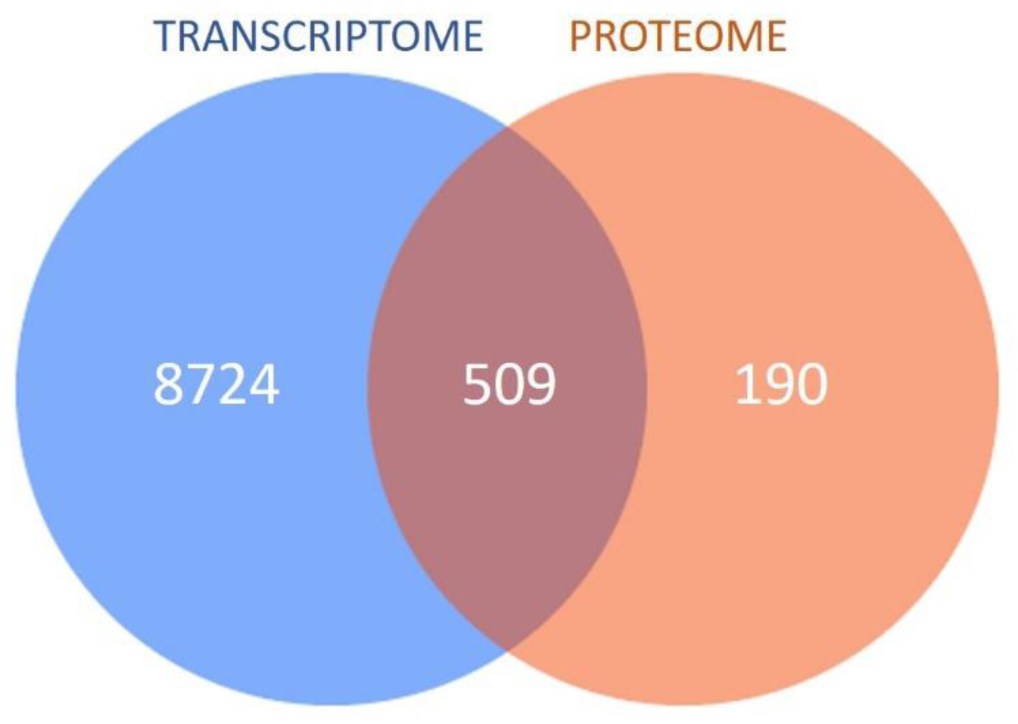
Venn diagram showing the number of transcriptome-specific, proteome-specific and conspecific genes and associated proteins identified during the transcriptomic and proteomic profiling of high-quality pikeperch eggs (n=8).

The set of transcriptome-specific genes could be organized into 5 Gene Ontology clusters (sharing at least 30% of genes) of the 30 most enriched biological processes, namely, neurogenesis, cellular component biogenesis, localization and transport of proteins, protein phosphorylation and transcription (Fig. 6a). KEGG pathway enrichment identified metabolic pathways, endocytosis and axon guidance, as the most enriched among the transcriptome-specific gene sets. In addition, it is noteworthy that protein processing in the endoplasmic reticulum was found among the top 5 most enriched KEGG pathways (Fig. 6d). Detailed Gene Ontology analysis of the groups of genes constituting each cluster confirmed their relevance to the processes in which they were grouped. The gene sets constituting particular clusters appeared to be related to the endocytosis, axon guidance, focal adhesion and MAPK signaling KEGG pathways (Supplementary file 8).

**Fig. 6.**
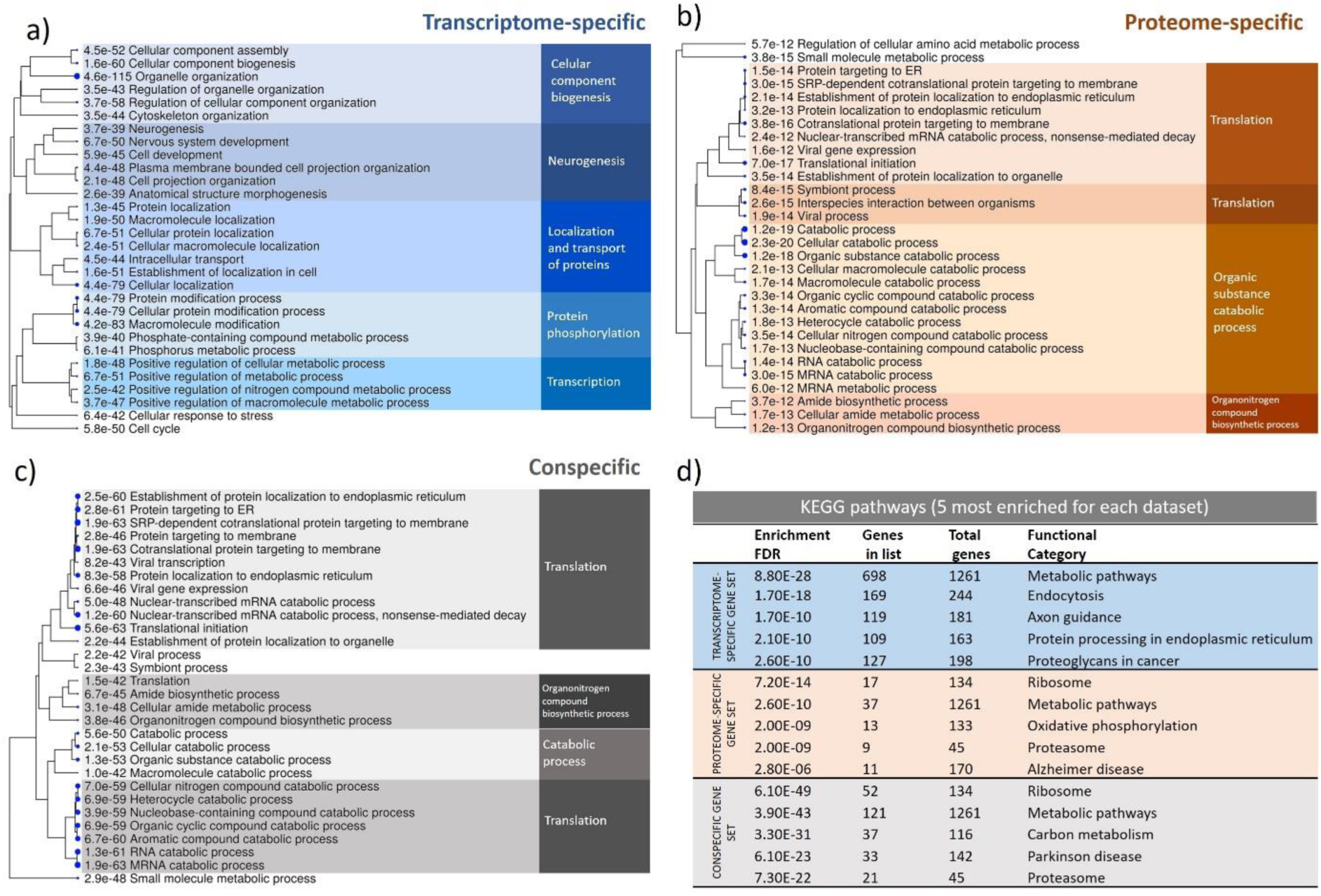
Clustering analysis of the 30 most enriched (FDR<0.05) biological processes (BPs) for (a) transcriptome-specific, (b) proteome-specific and (c) conspecific sets of genes and associated proteins identified during transcriptomic and proteomic profiling of high-quality pikeperch eggs (n=8). Entities constituting each cluster (grouped whenever the terms shared at least 30% of the genes/proteins) were again subjected to enrichment analysis to identify the main BP (for further details, see also Supplementary file 8). Additionally, the five most enriched KEGG pathways for each subset of data are presented in panel ‘d’.

Gene Ontology analysis of the proteins identified in pikeperch eggs revealed 4 different clusters comprising 28 out of the 30 most enriched biological processes (Fig. 6b). The data set analyzed here contained hits related to the ribosome and proteasome among the top 5 most enriched KEGG pathways. In addition, it should be highlighted that the proteome-specific gene set contained proteins related to metabolic pathways (similar to transcriptome-specific pathways) as well as protein modification activities, though oxidative phosphorylation appeared to be the primary pathway enriched (Fig. 6d). Furthermore, detailed Gene Ontology analysis of proteins constituting particular clusters indicated that the first two clusters were actually mostly related to translational processes. The proteins constituting the third cluster, comprising 13 Gene Ontology terms, were found to be related to various catabolic processes, including catabolism of RNAs and proteins. Proteins identified in the fourth and final cluster were found to be involved in the biosynthesis of organonitrogen compounds, with the biosynthesis of amides presumed to be primarily targeted in this process. KEGG pathway analysis of the genes constituting different clusters showed that the proteome-specific gene set was mostly related to the ribosome pathway, regardless of the cluster analyzed. In addition, it should be highlighted that the proteasome was among the most enriched KEGG pathways (Supplementary file 8).

Functional analysis of the gene set shared between the transcriptome and proteome allowed us to distinguish four clusters in which 27 out of 30 most enriched Gene Ontology terms were grouped (Fig. 6c). The genes in this gene set were found to be involved in similar KEGG pathways to those identified for the proteome-specific pathway. This includes ribosome, proteasome and metabolic pathways accompanied by carbon metabolism, among others (Fig. 6d). Specific functional analysis of the gene set constituting each of the Gene Ontology clusters revealed similarities with the analysis of the proteome-specific set of genes. Two clusters of genes were consistent and mainly involved in the translation process. The remaining clusters comprised genes involved in catabolic processes and organonitrogen compound biosynthetic processes. KEGG analysis revealed that in each cluster, the most enriched pathway was ribosomes, followed by carbon metabolism (in three clusters), proteasomes (in two clusters) and others, including RNA transport and metabolic pathways (Supplementary file 8).

### 3.2. Scientific objective no. 2: Transcriptomic-proteomic profiling of egg quality

#### 3.2.1. Evaluation of egg quality

Among the eggs exhibiting formation of the blastodiscs at the animal pole within 1 h post fertilization, two groups could be distinguished, i.e., eggs of high and low quality. Significant differences in embryonic survival could be detected at 72 h post fertilization (after ZGA). In the high-quality group, over 70% of larvae hatched, whereas in the low-quality group, less than 20% hatched. In both groups, less than 10% of hatched larvae were found to exhibit developmental deformities (Fig. 7).

**Fig. 7.**
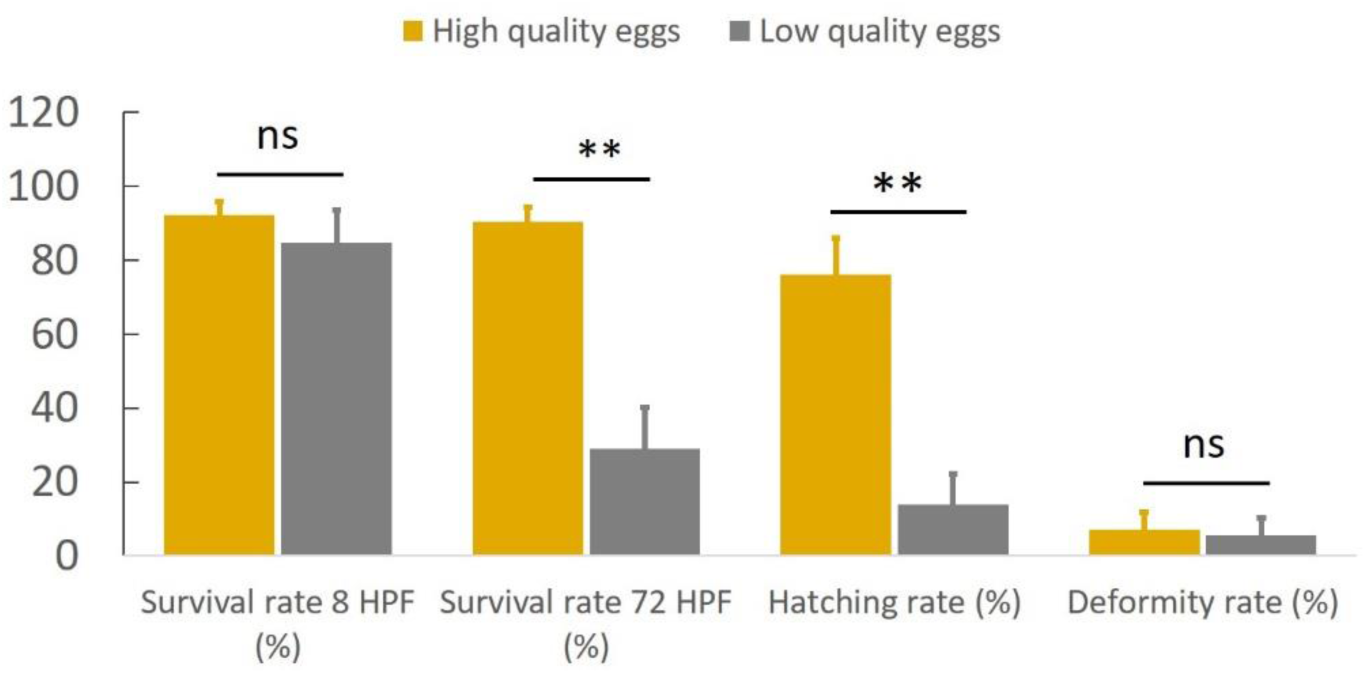
Results of evaluation of developmental competence of the preselected eggs assigned to high and low egg quality groups (n=5 for each group). ‘HPF’ stands for ‘hours post fertilization’; ‘ns’ stands for ‘nonsignificant’ (highlighting that no significant differences were detected; P>0.05). Double asterisks indicate that the data between the groups were significantly different (P<0.01).

#### 3.2.2. Transcriptomic analysis of high- and low-quality eggs

Within the comparative analysis of the transcriptomes of high- and low-quality eggs, successful hybridization was recorded for 17,243 probes. Statistical analysis revealed 84 differentially expressed genes (DEGs) (FDR<0.01 with at least a 2-fold change) (Fig. 8a, Supplementary file 9). Only 2 DEGs were found to be downregulated in high-quality eggs (*ctnn2b*, *msn*), and the remaining 83 DEGs were downregulated in low-quality eggs. Gene Ontology analysis revealed four different clusters (with at least 3 terms and sharing at least 50% of genes) that were found to be related to RNA splicing (two clusters), chromosome organization and cell junction organization. The most enriched term – organelle organization, associated with 31 DEGs – was not connected to any cluster (Fig. 8b).

**Fig. 8.**
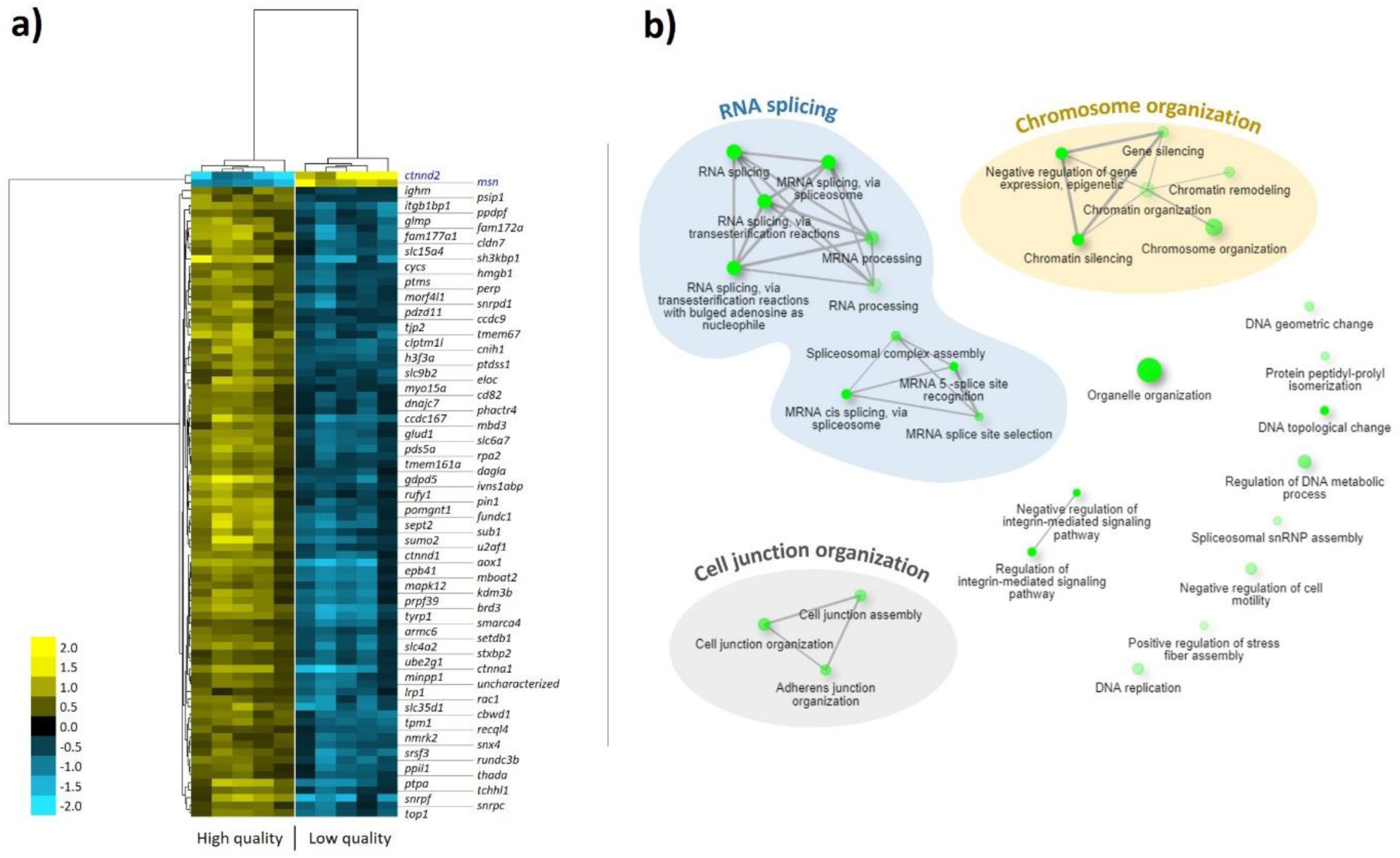
Panel ‘a’: Unsupervised average linkage clustering of 85 differentially expressed genes (DEGs) between high- and low-quality pikeperch eggs (n=5 for each group). Each row represents the same gene, whereas each column represents an RNA sample obtained from a batch of freshly ovulated eggs. The expression level for each gene is presented using a color intensity scale, where yellow and blue represent over- and under-expression, respectively. Black represents the median gene abundance. Panel ‘b’: Clustering analysis of the 30 most enriched ontological terms (darker nodes are more significantly enriched gene sets; larger nodes represent larger gene sets; thicker edges represent more overlapping genes). Genes in clusters highlighted with different colors were additionally verified to be involved in RNA splicing, chromosome organization and cell junction organization in a separate analysis.

By analyzing the expression levels of DEGs during the embryonic development of zebrafish, it was found that the expression levels of only 34 DEGs increased after ZGA. The remaining 47 DEGs were characterized by lower expression following ZGA (Fig. 9). qPCR validation of the 20 DEGs characterized by the highest expression before ZGA in zebrafish confirmed a significant difference (t-test, p<0.05) for 85% of them (Fig. 10).

**Fig. 9.**
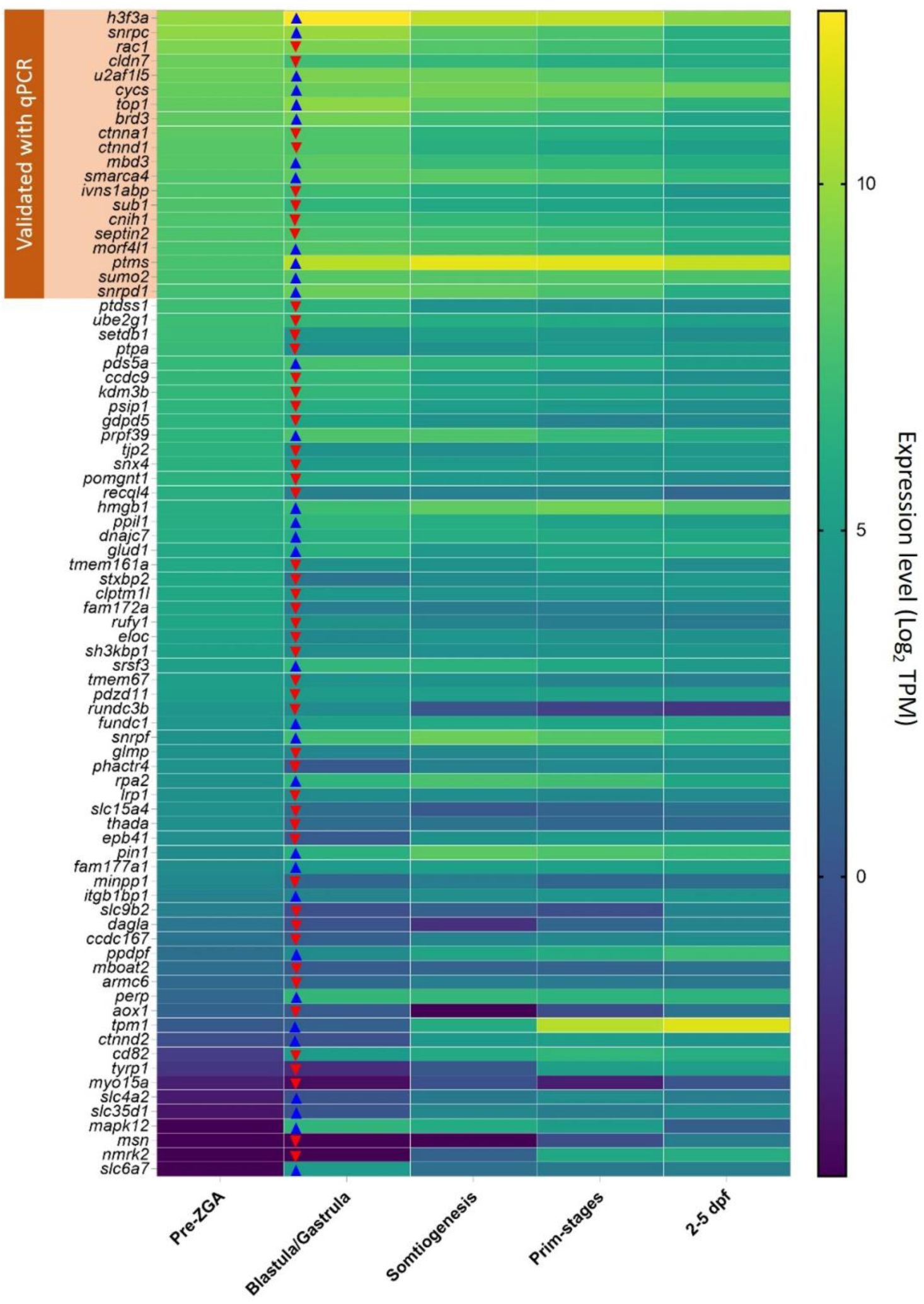
Heat map presenting the expression levels of differentially expressed genes identified during comparative transcriptomic analysis of high- and low-quality pikeperch eggs during zebrafish embryonic and larval development (based on data from White et al. 2017). Genes are arranged from top to bottom in descending order of the expression level based on the data provided in the first column (pre-ZGA; period before zygotic genome activation [ZGA]). Blue (up) and red (down) arrowheads provided on the blastula/gastrula stage column indicate up- and downregulation of particular genes following ZGA, respectively. TPM – number of transcripts per million reads, dpf – days post fertilization (2-5 dpf refers to the post hatching period). The first 20 genes (exhibiting the highest expression in the pre-ZGA period in zebrafish), highlighted and labeled in the upper-left corner, were subjected to qPCR validation of their expression level in pikeperch eggs (of various quality).

**Fig. 10.**
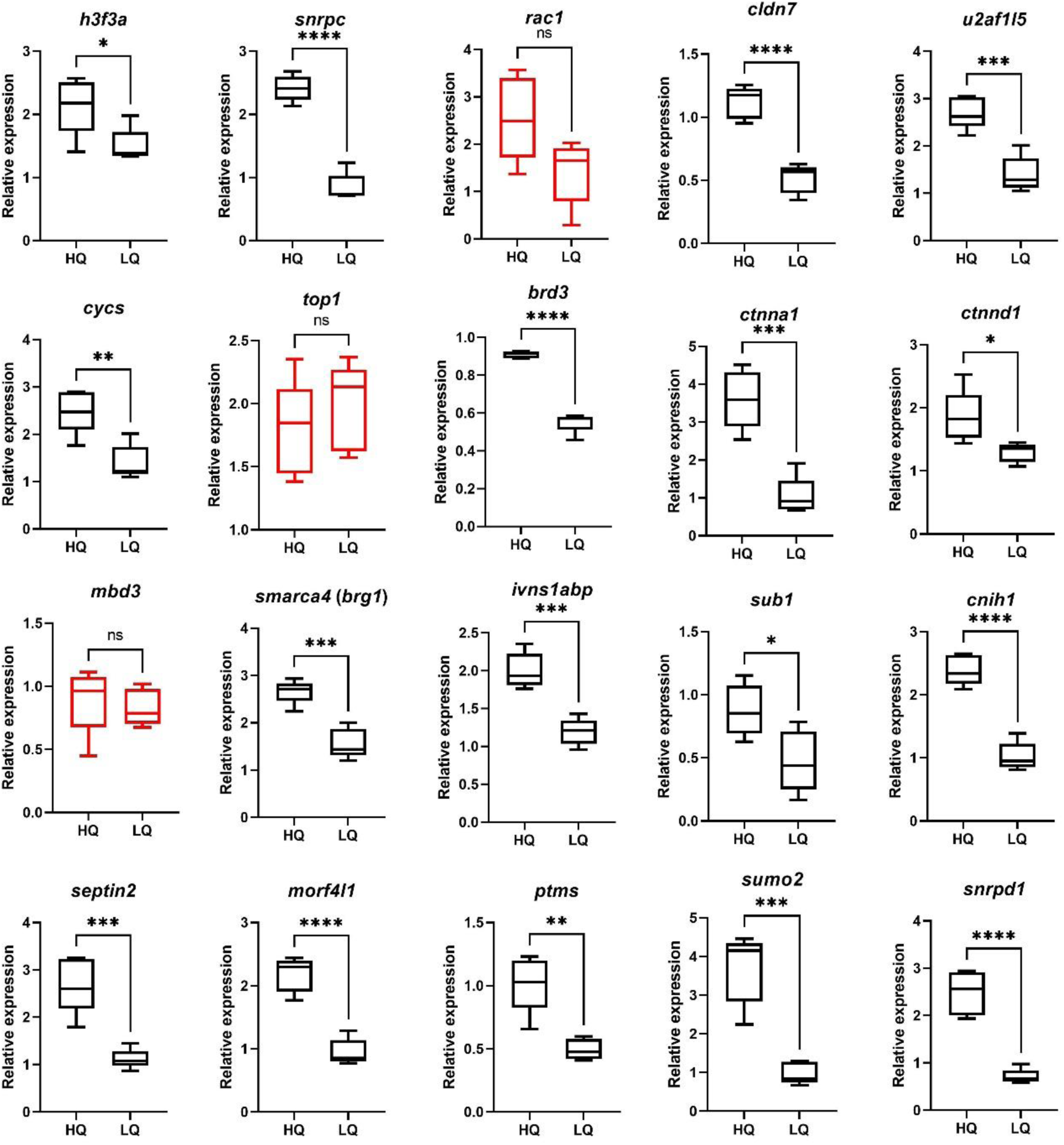
Results of validation of candidate genes (differentially expressed between high and low egg quality in pikeperch; n=5 for each group) with qPCR. Boxes represent the upper and lower quartiles, with the median value represented by the horizontal bar inside the box. Whiskers represent the minimum and maximum values recorded. The results of statistical analysis are presented as follows: ns – nonsignificant, * – P<0.05, ** – P<0.01, *** – P<0.001, **** – P<0.0001.

#### 3.2.3. Proteomic analysis of high- and low-quality eggs

Proteomic analysis of eggs of different quality enabled the identification of 946 proteins. Statistical analysis did not reveal any proteins with significantly different abundance between high- and low-quality eggs (FDR>0.05).

## 4. Discussion

### 4.1. Scientific objective no. 1

#### 4.1.1. Molecular profile of high-quality eggs

Our study provides novel insights into the understanding of the role of maternally-derived molecular cargo in fish. The two data sets contributed similarly to the highest level GO profile (Fig. 2). Both groups were associated with many processes that are critical for reproduction, development and the immune system. The same processes, while being very general, were already reported to be important components of the developmental competence of eggs in finfishes [17,21,59]. However, one unique feature of the transcriptome in this analysis that sheds light on the regulation of behavior should be mentioned. In a recent study, Żarski et al. (2020) indicated that genes involved in neurogenesis are highly modulated by domestication levels in the eggs of pikeperch. This suggests that the predetermination of nervous system development by maternally-inherited mRNAs should receive more attention. In particular, maternally-inherited mRNAs would shape embryo neurodevelopment and possibly the future behavior of the fish [60]. In contrast, in regard to proteomic-specific high level GO categories, the acrosome reaction category attracts attention, as it is well known that spermatozoa of teleost fishes have no acrosome [61]. This category included zona pellucida proteins, which are important components of egg envelopes that play various roles during oogenesis, fertilization and embryonic development [62]. It is worth mentioning that various forms of zona pellucida proteins were identified in the pikeperch egg proteome (Supplementary file 4), confirming their importance in the cells analyzed.

Further analysis shows, for the first time, that transcripts and proteins have distinct, yet complementary, functions in the egg of teleost fish. The clustering analysis of most enriched GO terms indicates certain unique functionalities for each of the two data sets analyzed. It should be highlighted that among the specific GO terms, those involved in neurogenesis were characteristic of the transcriptome (Fig. 3, Supplementary file 7). On the other hand, high representation of molecules responsible for the immune response could be identified as characteristically enriched in the proteome and was not found in the top 500 enriched processes for the transcriptome data set (Supplementary file 4). Obviously, immune response genes are important mRNAs identified in eggs of percids [63] and in pikeperch (Supplementary file 3). However, given that translation is quiescent, it seems to be more likely that the proteins already present in the egg play a main role in the defense against pathogens in eggs and early embryos. Considering that several important immune response translation proteins are among the most abundant proteins (next to vitellogenins and zona pellucida proteins), including galactose-specific lectin nattectin-like proteins [64], it is clear that defense against pathogens is among the priorities for fish eggs. It was previously suggested that immune response-dependent protein profile in eggs can be modulated by the rearing environment in which females are kept [48]. Therefore, it should be highlighted that health and welfare status of broodfish before FOM, during which the majority of proteins are deposited [12], may affect the capacity of the eggs and developing embryo to tackle the pathogens. However, the role of the transcriptome in defense mechanisms should not be neglected, because genes encoding immune response proteins are also present in the egg at very high levels, including the galactose-specific lectin nattectin protein (third most expressed gene). This indicates that this protein may play a crucial role throughout embryogenesis as a first line of immune defense. However, further, more dedicated study of this issue is required to draw valid conclusions.

#### 4.1.2. Role of transcriptome-specific, protein-specific and conspecific molecules

Molecular profiling revealed characteristic features of the transcriptome-specific subset of genes. The mRNA contained in the oocyte was found to be considerably enriched in processes related to transcription, protein phosphorylation, biogenesis of cellular compounds and localization and transport of the proteins. This highlights that maternally-inherited transcriptomic cargo plays a crucial role in the intracellular ‘management’ of the proteins produced but is also responsible for gene expression (features also recorded at the level of the whole transcriptome; see Fig. 3). However, the most unique characteristics were associated with the genes involved in neurogenesis; this was additionally confirmed by KEGG pathway enrichment analysis, which identified axon guidance as among the most enriched pathways (for details see Supplementary file 8). It should be emphasized that axon guidance was already reported to be modified by the domestication process in pikeperch eggs [47]. Therefore, our study clearly indicates that development of the nervous system is a characteristic ‘maternal legacy’ in the ovulated egg and possibly in the developing embryo as well. This again brings attention to behavioral traits as factors conditioning adaptability of the fish to particular habitats and/or culture conditions.

Functional analysis of proteome-specific and conspecific subsets of data for the two types of molecules indicates the high importance of translation under the control of both types of molecules. This is in accordance with previous reports indicating that translation is among the crucial processes conditioning developmental competence in fishes [1, 21], similar to the case in other animals [18]. In addition, clustering revealed the importance of protein and nucleic acid catabolism as well as organonitrogen compound biosynthesis, for which the cell is prepared at both the transcriptomic and proteomic levels. Interestingly, the KEGG pathway enrichment analysis highlights, along with metabolism or oxidative phosphorylation, two major elements: the ribosome and the proteasome (see also Supplementary file 8). This additionally confirms the very great importance of intracellular biosynthesis of proteins and their degradation, suggesting extensive remodeling within the eggs. Żarski et al. [21] reported that genes involved in ubiquitination (intracellular protein degradation) are crucial for the developing embryo. Additionally, Chapman et al. [23] reported that the expression level of ubiquitin-related genes in ovaries was related to developmental competence of eggs in striped bass (*Morone saxatilis*). Our study confirms that the potential for translation and degradation of proteins and nucleic acids is among the most important features of ovulated fish eggs.

### 4.2. Scientific objective no. 2

#### 4.2.1. Molecular profiling of egg quality

Egg quality recorded during the controlled reproduction of pikeperch is usually highly variable [35, 38]. In fish, such variability is derived from both unfertilizable eggs (e.g., overripened eggs) and eggs that are competent for fertilization but die during the early stages of embryonic development [17]. The latter phenomenon, still barely studied in fish, could be considered in our study thanks to the specific preselection procedure applied (Fig. 1). This allowed us to find, for the first time in percids, that such early embryonic lethality significantly contributes to the overall egg quality. However, it should be noted that the eggs exhibiting high early embryo lethality did not contribute to an observed increase in the incidence of larval deformity, which is a growing and serious concern in percids [46]. Therefore, the eggs preselected for molecular analysis in our study were characterized by high fertilization capacity but different developmental potential following fertilization. Thus, we were able to focus on the molecular profile being predictive of altered ‘machinery’ leading to developmental failure at early stages, which was done for the first time with such complex molecular profiling.

Among the two types of molecules investigated in our study, only the transcriptomic profile was found to be predictive of egg quality, whereas proteomic profiling did not reveal any significant differences between high- and low-quality eggs. This is in contrast to other studies revealing differentially abundant proteins in eggs of different quality. For example, in Eurasian perch [26] and in zebrafish [25], respectively, tens and hundreds of differentially abundant egg-quality-dependent proteins were detected. Such discrepancies between the published studies and the data obtained in our study may arise for various reasons. In the case of Eurasian perch reported by Castets et al. [26], it can be presumed that the sampling strategy, without any preselection, could affect the overall proteomic profile observed. However, in the case of zebrafish, almost all the eggs analyzed developed to the approximately 256-cell stage, indicating that embryonic development failed after successful fertilization. However, the main reason for the discrepancies observed could be associated with the different proteomic approaches that were used. In the study of Yilmaz et al. [25], proteomic profiling was performed after initial molecular weight-based fractionation of the proteins, which increased the number of quantified proteins. However, in the case of zebrafish, highly abundant proteins (such as vitellogenins, zona pellucida proteins or ribosomal proteins) were also found to be differentially abundant between high- and low-quality eggs. Therefore, it could be assumed that our straightforward LC-MS/MS approach without prefractionation allowed us to identify egg-quality-related proteins and should detect protein-dependent processes affecting egg quality. Of course, the discrepancies in overall findings between the studies may also be explained by species-specific factors and egg characteristics. Species specificities in the molecular profile of eggs have also been evidenced at the transcriptome level in a comparative analysis of cod and trout eggs [65]. Nevertheless, it is clear that we did not find strong egg quality-dependent proteome alterations in pikeperch, which could also be related to the applied preselection procedure. This may suggest that early embryo lethality in this species is determined after the generation of the protein reservoir in the egg (ending before the start of the FOM process; [12]), which highlights the fact that processes occurring during the FOM are strong modulators of egg quality. This finding is in agreement with a recent study suggesting that the processes directly preceding ovulation, and not those occurring during oogenesis, are the main modulators of egg quality [17]. However, to draw final conclusions, more detailed research is still needed.

#### 4.2.2. Candidate markers of egg quality

Among 85 DEGs identified to be predictive of early embryo lethality in our study, only 2 (*ctnnd2* and *msn*) were downregulated in high-quality eggs, whereas the remaining genes were downregulated in low-quality eggs (Fig. 8). The identified DEGs were found to be enriched in organelle organization, RNA splicing, chromosome organization and cell junction organization. Interestingly, none of these ontology terms have been previously reported to be clearly related to egg quality in teleosts. However, when mapping these DEGs to the processes enriched in the whole transcriptome (see Fig. 3) and proteome (Fig. 4), it becomes apparent that this set of genes plays multiple roles in the cell, including in protein modification, transcription, and translation and in cellular signaling, which were previously reported to have egg-quality-modulatory effects in other species [17,21,24,59,66]. Interestingly, a high number of DEGs (33) were also found to play a role in neurogenesis, representing the first time that egg quality markers have been linked with this important biological process. These observations highlight the added value of the combined ontological analysis (typical enrichment analysis and global overview of the whole transcriptome) implemented here, as it provides new insights into the roles of DEGs in the developmental competence of pikeperch eggs and in our study draws attention to the multifunctionality of the gene set. Moreover, it highlights particular genes as modulators of developmental competence rather than involved solely in the processes for which this gene set is enriched. From this point of view, our study provides novel insights into the processes determining egg quality.

A comparison of published data with the set of DEGs identified in the present study indicates the uniqueness of the data obtained since none of the candidate genes obtained for pikeperch was reported to be quality-dependent in various fish species [1,17,21,22,67,68]. Therefore, we examined at the expression profile of all the DEGs during embryogenesis in zebrafish based on data published by White et al. [56]. This allowed us to find that not all the genes were expressed at the zygote stage in zebrafish, despite being relatively well expressed in pikeperch eggs (see Supplementary file 3). This includes *slc4a2*, *slc35d1*, *mapk12*, *msn*, *nmrk2* and *slc6a7* (Fig. 8). This analysis reveals that the maternal transcriptomic cargo contained in the eggs may vary among species. A similar conclusion was drawn by Myers et al. [69], who reported that genes constituting quality markers in other species (as reported by Sullivan et al. [66]) were also poorly expressed or not at all expressed in ovulated eggs of hybrid catfish until neurulation. Interspecies differences in transcript abundance in eggs between cod and rainbow trout were also reported by Wargelius et al. [65]. This pattern highlights that the interspecies comparison of the molecular markers related to egg quality may suffer from very large biases. It also allows us to suggest that the molecular mechanisms driving early embryogenesis may also vary across species. However, this needs to be studied in greater detail, with similar analytical workflows for data collection (laboratory techniques, bioinformatics, etc.) to draw valid conclusions. Nevertheless, our study is the first to provide a set of molecular markers of early embryo lethality in pikeperch, which can be a valuable tool in further studies on the developmental competence of eggs in this important fish species.

The selection of candidate genes based on expression patterns during zebrafish embryonic development led to the identification of 20 genes that are crucial for embryonic development (Fig. 9). Importantly, 17 of them (85%) were successfully validated with qPCR, providing a set of solid candidate quality markers crucial for development in fish and other animals. For example, *h3f3a* was found to be an essential maternal factor for successful oocyte reprogramming in mice [70], and *snrpc* was previously reported to play a crucial role during amphibian and fish development [71]; *cldn7* was responsible for maintaining blastocyst development in pigs [72], whereas *ctnna1* was reported to be important for the development of the heart during mammalian embryogenesis [73]. Moreover, *ctnnd1* expression, which is dependent on the level of luteinizing hormone, was found to be crucial during maturation in human oocytes [74], suggesting that this gene could be a valuable marker of not only egg quality but also FOM progression. Another gene, *smarca4* (also known as *brg1*), was reported to regulate ZGA in the mouse [75], which makes it a highly promising candidate gene for monitoring early embryonic lethality stemming from failure during ZGA in fishes. Several identified genes are important in mammals for processes related to the progression of oogenesis (*sub1*; [76]), oocyte development (*sumo2*; [77]) and meiosis (*septin2*; [78]), further highlighting the possibility of using these markers in developmental studies in fishes. In the case of the remaining positively validated transcripts, such as *u2af1l5*, *cycs*, *brd3*, *ivns1abp*, *cnih1*, *morf4l1* and *ptms*, their involvement in early embryogenesis remains to be explored in fishes and other animals, as they likely constitute a valuable set of candidate genes.

### 4.3. Conclusions

Our study provides novel insight into the role of maternally-inherited molecular cargo in finfishes. The approach taken in our study sheds light on the importance of the transcriptome in the development of the nervous system, which confirms the role of neurogenesis-related mRNAs as very important nongenetic heritable factors [7, 79]. On the other hand, proteomic analysis highlights the crucial and specific role of proteins in the immune response in ovulated eggs. Additionally, integrated transcriptomic-proteomic analysis draws attention to the galactose-specific lectin nattectin gene and protein as the frontline defense molecule for the egg and – most likely – the developing embryo.

The molecular analysis of egg developmental competence emphasizes postvitellogenic processes (FOM and ovulation) as those compromising the transcriptomic profile but not affecting proteomic cargo. This brings attention to the need for careful reconsideration of the prespawning conditions that the fish are exposed to and reproductive protocols (previously reported to affect egg quality [38] as well as transcriptomic profile [22]) as very strong modulators of egg quality and their molecular structure. Although the mechanisms driving these alterations and the consequences stemming from the differential abundance of the transcripts remain to be explored, the candidate quality markers provided in this study are the first to be identified in pikeperch and represent valuable resources for further studies on the reproductive biology of this and other fish species. All of these results are of major importance for understanding the influence of external factors on reproductive fitness in both captive and wild-type fish species.

## Acknowledgements

The authors are sincerely grateful to ASIALOR fish farm for their support and access to their facilities as well as Jean Babtiste Muliloto, Miroslav Blecha and Maud Alix for their assistance during the reproduction of pikeperch.

## 6. Data availability statement

Raw data from the comparative analysis of eggs of high and low quality can be accessed via the NCBI Gene Expression Omnibus under the GSE167376 accession number [*the data will be made publicly accessible following publication*].

The mass spectrometry proteomics data have been deposited in the ProteomeXchange Consortium via the PRIDE partner repository with the dataset identifier PXD023229 [*the data will become freely accessible following publication*].

## 8. Contribution of the authors

**Daniel Żarski**: conceptualization, methodology, investigation, validation, formal analysis, visualization, supervision, project administration, funding acquisition, writing – original draft

**Aurelie Le-Cam**: validation, formal analysis

**Thomas Frohlich**: validation, formal analysis

**Miwako Kösters**: investigation

**Joanna Nynca**: investigation

**Chrostophe Klopp**: data curation

**Sławomir Ciesielski**: investigation

**Beata Sarosiek**: investigation

**Katarzyna Dryl**: investigation

**Jerome Montfort**: data curation, validation

**Jarosław Król**: investigation

**Pascal Fontaine**: resources, supervision

**Andrzej Ciereszko**: funding acquisition, supervision

**Julien Bobe**: conceptualization, methodology, supervision

## Supplementary information

**Supplementary file 1:** Schematic diagram illustrating the Gene Ontology analysis strategy applied during the functional analysis of the transcriptome and proteome obtained from high-quality pikeperch eggs.

**Supplementary file 2:** Primers used for qPCR validation of candidate genes for biomarkers of egg quality in pikeperch.

**Supplementary file 3:** List of genes identified in high-quality pikeperch eggs (n=8). For each entry, the gene name and accession number of the human homolog were provided, except for those identified as duplicates (when different genes mapped to the same protein) and fish-specific genes. In addition, the mean intensity signal (for n=8) was provided, and genes were arranged in descending order of signal intensity.

**Supplementary file 4:** List of all proteins identified in high-quality pikeperch eggs (n=8). For each entry accession number, the mean signal intensity and protein score (i.e., number of samples in which particular protein was present) are given. Proteins are presented in descending order based on signal intensity.

**Supplementary file 5:** List of abundant (present in at least 50% of the samples) proteins in high-quality pikeperch eggs (n=8). Each entry-associated gene name and accession number (for human homolog) was provided, except for the fish-specific proteins. In addition, the mean intensity signal (for n=8) was provided, and genes were arranged in descending order based on signal intensity.

**Supplementary file 6:** Higher level Gene Ontology categories (in alphabetic order) identified for whole transcriptome and proteome data obtained from high-quality pikeperch eggs (n=8). For each category, the numbers of genes and proteins identified in either the transcriptome or proteome are provided.

**Supplementary file 7:** Most enriched (top 500 biological processes) ontology terms obtained during transcriptomic and proteomic profiling of high-quality pikeperch eggs. The ontological terms were split into transcriptome-specific, proteome-specific and conspecific groups.

**Supplementary file 8:** Results of the ontology analysis (enrichment analysis of biological processes and KEGG pathways) of transcriptome-specific, proteome-specific and conspecific subsets of genes/proteins obtained from high-quality pikeperch eggs based on clusters obtained in the analysis presented in Fig. 5.

**Supplementary file 9:** List of 85 differentially expressed genes (DEGs) identified in transcriptomic profiling of pikeperch eggs characterized by high (n=5) and low (n=5) quality. For each entry gene name, the protein name (encoded by the respective gene), accession number and human homolog (whenever available) used for functional analysis are presented. Additionally, zebrafish homologs used during the evaluation of expression levels during zebrafish development were provided whenever they could be identified. In addition, the fold change difference (between high and low egg quality) in expression was also provided. Genes are arranged alphabetically based on gene name.

