## Supplementary figures and images for "Neurodevelopment vs. the immune system: complementary contributions of maternally-inherited gene transcripts and proteins to successful egg development in fish"

### Supplementary file 1

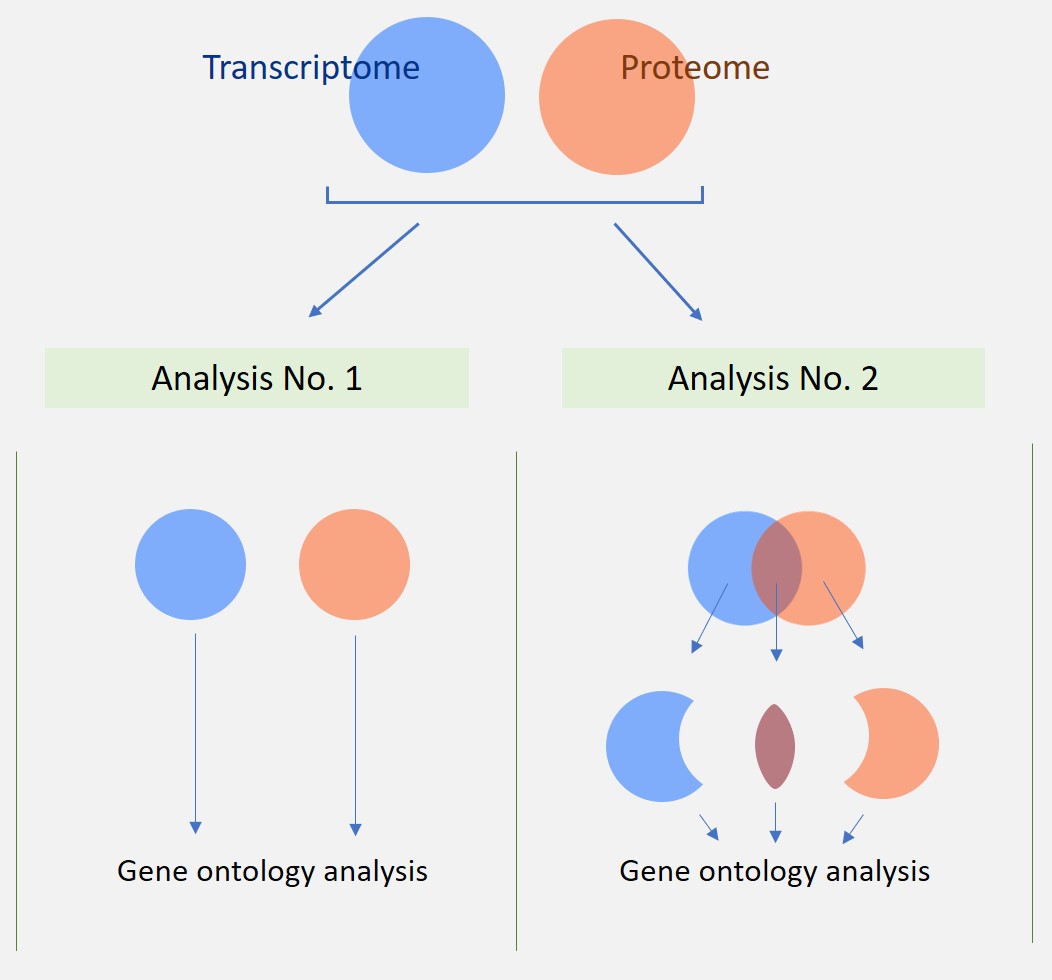
