## Supplementary file 2 for "Neurodevelopment vs. the immune system: complementary contributions of maternally-inherited gene transcripts and proteins to successful egg development in fish"

Primers used for qPCR validation of candidate genes for biomarkers of egg quality in *Sander lucioperca*.

| Gene symbol | Gene name | Forward primer | Reverse primer |
| --- | --- | --- | --- |
| Candidate genes | | | |
| *h3f3a* | histone H3-like centromeric protein CSE4 | GTCAAGAAGCCCCATCGTTA | TCCTCGAATAGACCCACCAG |
| *snrpc* | U1 small nuclear ribonucleoprotein C | CACATGATTCGCCATCAGTAA | GGGAGGAATCTTTCCCTGTT |
| *rac1* | ras-related C3 botulinum toxin substrate 1 | GTGATGGTTGATGGGAAACC | TCTCTCAGGTCCAGCTTGGT |
| *cldn7* | claudin-7-B | TCAATCCTGCAGCTCAACAG | TAATGCCACCAGTCATGGAA |
| *u2af1l5* | splicing factor U2AF 35 kDa subunit | ACTAAGTGGTCCCCGAGGTT | CGCTGCTCCAGATTACACAA |
| *cycs* | cytochrome c | TGCCTGCAGACATTGAAAAA | ATTCCAGACAATGCCTTTGC |
| *top1* | DNA topoisomerase I, mitochondrial-like | GATTGAGGAGGCAAGACTCG | GTCCAACATCTGGGCAAAGT |
| *brd3* | bromodomain-containing protein 3 isoform X1 | ATTCACACAACGCCACAAAA | GGTAATCGTGTAGCCCCAGA |
| *ctnna1* | catenin alpha-1 | TGAAGTAGGCGAGGAAAGGA | GAAAAACATCCCGCAGTTGT |
| *ctnnd1* | catenin delta-1 isoform X3 | GACAACACCTGCCAAAGGTT | GGGAAGACCCTTCTCCAGAC |
| *mbd3* | methyl-CpG-binding domain protein 3-like isoform X1 | GCTAGAGGAGGCACTGATGG | ACTGCTGTATGGCTCCGACT |
| *smarca4 (brg1)* | transcription activator BRG1 isoform X3 | TGGAGAAAGACGTGATGCTG | TCGTCGTCCTCTTCCTCACT |
| *ivns1abp* | influenza virus NS1A-binding protein | CCAGCTAAAGGCAGACAAGG | CTCCCATAGAGCTGGCAAAG |
| *sub1* | activated RNA polymerase II transcriptional coactivator p15 isoform X1 | CAAAGAGTGGCGAGAGTTCC | TCTCCCCATCTTGGTTCATC |
| *cnih1* | protein cornichon homolog 1 isoform X2 | GGCTCACCCTCTGTCTCAAC | TTTGCACCAGCCTTCTTTTT |
| *septin2* | septin-2B isoform X1 | TCAGGTTCATCGCAAGTCTG | GCGTCTCCATATCCTGGTGT |
| *morf4l1* | mortality factor 4-like protein 1 | CTTGTGGATGACTGGGACCT | GATACCAGCAACCACCTCGT |
| *ptms* | parathymosin-like | GGTCTCTGCCCTGCTACTTG | GATGCAGGGCCACTATGTTT |
| *sumo2* | small ubiquitin-related modifier 2-like isoform X2 | AAAGGCACACTCCTCTCAGC | TGTTGGAACACGTCAATCGT |
| *snrpd1* | small nuclear ribonucleoprotein Sm D1 | TGAAACTCAGCCATGAGACG | TTGTTGCCACGGATACTCAG |
| Normalizing genes | | | |
| *xrn1* | 5'-3' exoribonuclease 2 isoform X2 | ACGAACCGAACTTCACCATC | AGACACTGGCATCGTGTCTG |
| *cd81 antigen-like* | CD81 antigen-like isoform X1 | GCTGATTGTGGTGGTGTTTG | CCAATGACTGGGTCTCCACT |
| *col14a1* | collagen alpha-1(XIV) chain | CAGAGCTTGGCCTTCAAAAC | CCTAAACCAAAGGCCAGTGA |
| *ubql4* | ubiquilin-4-like | TGACGGTTCACCTGGTCATA | GTATGTTGGGCGTCTGTGTG |
